# Neural Ensembles in the Lateral Prefrontal Cortex Temporally Multiplex Task Features During Virtual Navigation

**DOI:** 10.1101/2024.01.10.574378

**Authors:** Mohamad Abbass, Benjamin Corrigan, Renée Johnston, Roberto Gulli, Adam Sachs, Jonathan C. Lau, Julio Martinez-Trujillo

## Abstract

Neuronal populations can expand their information encoding capacity using mixed selective neurons. This is particularly prominent in association areas such as the Lateral Prefrontal Cortex (LPFC) that integrate information from multiple sensory systems. During naturalistic conditions where subjects have agency, it is unclear how LPFC neuronal ensembles process space and time varying information. Here we show that during a virtual navigation task that requires associative memory and decision making, individual neurons and neuronal ensembles in the LPFC time-multiplex their selectivity for different task features to fluidly encode the temporal contingencies of the task and the animal’s choice. Neurons in ventral regions showed more selectivity for non-spatial features, while dorsal regions showed selectivity for space and eye movements. These results demonstrate that during naturalistic tasks with fluid scenery and spatiotemporal dynamics, LPFC neurons and neuronal ensembles time-multiplex the components of their selectivity, expanding the spatiotemporal capacity of their neural codes.

## Introduction

The primate lateral prefrontal cortex (LPFC) sits on top of the sensorimotor processing hierarchy and has been implicated in cognitive functions, including selective attention, working memory and rule encoding (Duncan, 2001; Miller & Cohen, 2001; Petrides, 2005a). Single neurons in the LPFC have been reportedly tuned to multiple task features in visuomotor tasks (Asaad et al., 1998; Mansouri et al., 2020; Wallis et al., 2001). This property has been termed mixed selectivity, which can be linear or non-linear, depending on how feature components are ‘mixed’ in neuronal firing rates (Rigotti et al., 2013). Mixed selectivity has been suggested to be behaviorally and computationally relevant, as it can increase the dimensionality of representations and facilitate read-outs of task-relevant features (Parthasarathy et al., 2017; Rigotti et al., 2013). However, most tasks exploring selectivity in LPFC neurons have been conducted using simple visual displays and constraining eye movements. It remains unclear how individual neurons and neuronal ensembles in LPFC mix information in scenarios with complex dynamic scenery, unconstrained eye movements and sensorimotor events occurring in a continuous and time-fluid manner.

The LPFC is heavily interconnected with numerous cortical areas and has been cytoarchitecturally divided into areas 8, 9, 10, 12, 45 and 46v/d, with tracing studies describing distinct anatomical connectivity patterns across these areas (Barbas, 2015; Barbas & Pandya, 1989; Petrides, 2005b). A recent study revealed distinct functional connectivity patterns in the dorsal (e.g., areas 9/46) and ventral (areas 45/47) LPFC, corresponding to dorsal and ventral high-level sensory areas (Xu et al., 2022). However, a functional organization to the LPFC has not been as clearly delineated. Single neurons in the dorsal and ventral LPFC have been reported to preferentially represent spatial and visual/object information respectively (Meyer et al., 2011; O’Reilly, 2010; Riley et al., 2016). However, representations of numerous modalities and levels of sensory integration have been reported in both dorsal and ventral LPFC (Kadohisa et al., 2015; Siegel et al., 2015; Tang et al., 2021). These findings may be linked to the integrative nature of the LPFC, with its ability to merge task-relevant information (Duncan, 2001; Miller & Cohen, 2001).

Previous studies have shown that neurons in the LPFC can become selective for combinations of relevant features during associative learning tasks (Mansouri et al., 2020; Rouzitalab et al., 2023; Wallis et al., 2001). However, these studies have used behavioural tasks involving controlled stimulus presentation of few task-relevant features in simple displays while constraining eye movements. The LPFC must be able to perform its function in real-world conditions, while subjects perform complex tasks and explore visual scenes via gaze shifts. Tremblay et al. (2022) demonstrated that single neurons in the LPFC maintain their representation of task features during unrestrained gaze movements. Moreover, Roussy et al. (2021) showed that representations of spatial locations by many LPFC neurons during virtual reality navigation in complex visual environments remain robust to changes in gaze position. These studies suggest that many LPFC neurons contain spatiotopic representations of the environment. More recently, Corrigan et al. (2023) have shown that LPFC neurons encode views of visual scenes suggesting that LPFC neurons may mix spatial and non-spatial information. However, it is unclear whether these components of mixed selectivity multiplex over time or are mixed together in a single multidimensional representation.

Here, we investigate how mixed selective neurons in LPFC (areas 9/46) encode spatial and non-spatial features of the environment during an associative memory task that requires virtual navigation through a complex visual environment. To this end, we recorded the responses of hundreds of neurons in two monkeys (*Macaca mulatta*) as they completed the task and freely explored the different elements of the visual scene via gaze shifts. Importantly, the task had different periods in which different features serially appeared. We found that individual neurons and neuronal ensembles in the LPFC time-multiplexed their representations of task relevant features and space. The dorsal and ventral LPFC had different tuning profiles and temporal dynamics. The ventral LPFC preferentially represented non-spatial task-features whereas the dorsal LPFC predominantly encoded space.

## Results

We trained two male monkeys (*Macaca mulatta*) to perform a context-color association task in a virtual environment (**Figure 1**). The animals used a joystick to navigate in an X-shaped maze, with each trial beginning in one arm of the maze (**Figure 1c**). Once the monkeys entered the corridor the walls changed texture to steel or wood (the context). When the monkeys reached the end of the corridor two coloured discs appeared, one at the end of each arm of the maze. The location of the colors (left or right arm) was randomly determined for each trial (color pair order or CPO). The context determined the color disc which the monkeys needed to navigate to for a reward, termed target side (**Figure 1d)**. Monkeys performed a fixed association trial block and then learned new associations each day. Performance was significantly above the 50% chance level (**Figure 1e** and methods). Across all sessions, Monkey B had a mean (± standard error) of 80.9 ± 0.02% correct trials, and monkey T had a mean of 70.3 ± 0.03% correct trials.

**Figure 1.**
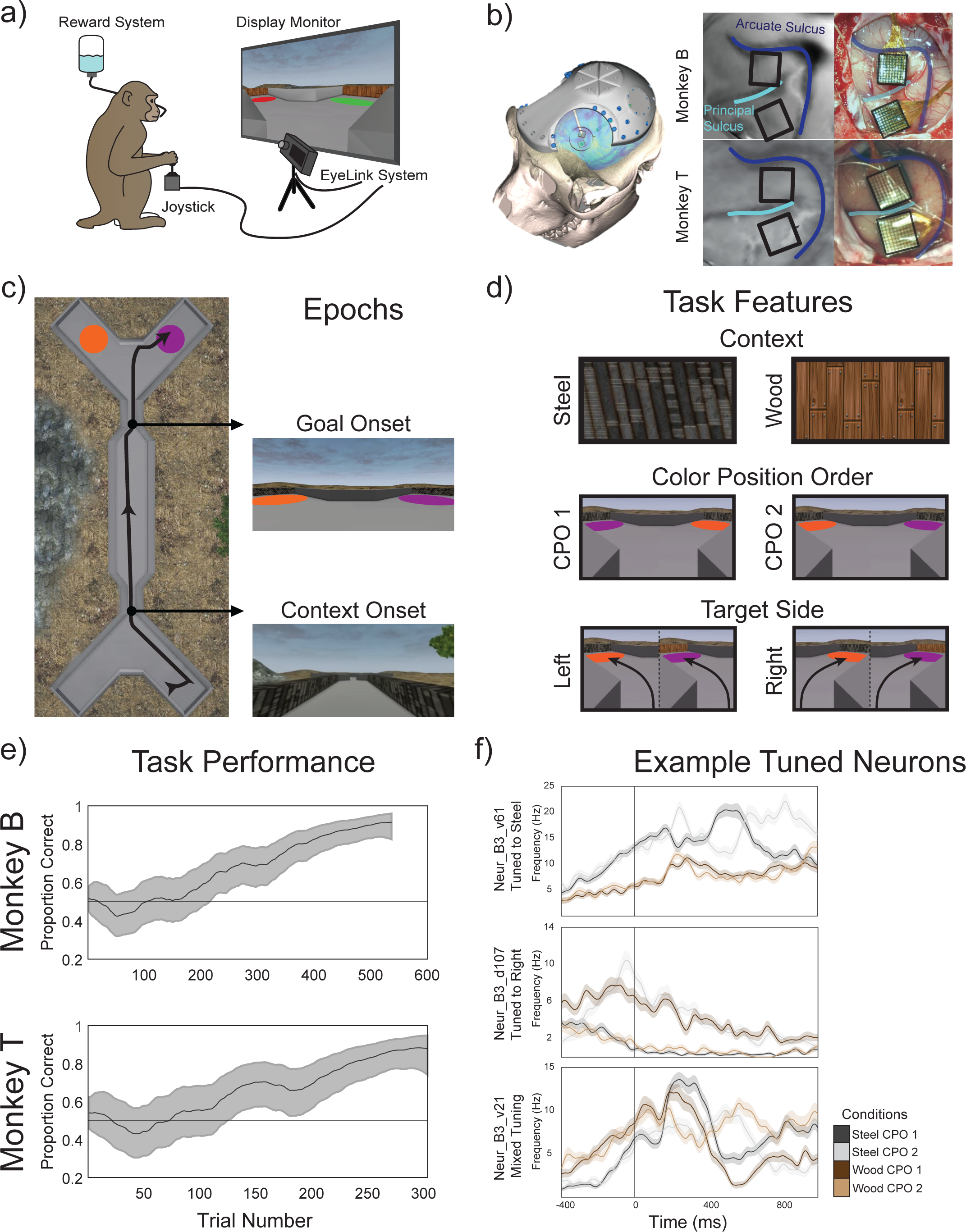
Recordings from NPH LPFC during a virtual navigation associative memory task. a) Monkeys were seated in front of a monitor and used a joystick to navigate through the virtual reality maze to complete the associative memory task. b) Location of implanted Utah microelectrode arrays, placed in the dorsal (areas 9/46d) and ventral (area 9/46v) lateral prefrontal cortex c) Overview of X-maze, with an example path taken during one trial, with the onset of the context (walls changing to steel or wood) and goal (two possible colors per session appearing in a random order) indicated. d) All possible task conditions, with context, color pair order (CPO) and target side. e) Estimated learning state in both monkeys with 95% confidence interval in example sessions with a novel context-association. f) Mean firing rate over 400ms in example neurons tuned to task features during goal onset, tuned to steel (above), right (middle) and mixed selectivity (bottom).

### LPFC neurons are tuned to task features during virtual navigation

While the monkeys completed the virtual reality navigation task, we recorded from a total of 813 neurons (512 in monkey B and 301 in monkey T) during six sessions. Single neurons were isolated manually (see methods). We computed firing rates and spike density functions for each isolated neuron and condition and found many neurons gave distinctive responses between conditions (see example **Figure 1f**). Single neuron tuning to task-related variables was determined using a multivariate linear regression model (see methods). Using correct trials, we investigated the proportion of neurons tuned to context, CPO, and target side. In this case, mixed selectivity refers to selectivity to two or more of the above variables. We investigated the proportion of tuned neurons over a 1200ms window (-200ms to 1000ms) around the context and goal onset epochs. During the context onset, we found significantly more neurons tuned to context in the ventral LPFC compared to the dorsal LFPC: 8.3% compared to 1.3% in monkey B (Chi-Square, χ = 16.71, p<0.001) and 9.3% compared to 2.5% in monkey T (Chi-Square, χ = 5.39, p<0.05). Here ventral and dorsal LPFC represent the location of the MEA relative to the principal sulcus. The dorsal array covers regions of areas 8Ad and 46d. The ventral array covers the ventral subdivisions of these areas, 8Av and 46v, and may extend to area 45 (estimation according to Petrides, 2005).

During goal onset, the appearance of the colors allowed the monkeys to determine which side to navigate towards. During this epoch, both monkeys had neurons in the ventral LPFC tuned to context (12.4%), CPO (6.2%), target side (17.2%), and demonstrated mixed selectivity (21.2%; **Figure 2**). Additionally, both monkeys had a significant number of neurons in the dorsal LPFC tuned to target side (27.2%); however, only monkey B had a significant number of neurons with mixed selectivity (19.5%). The dorsal and ventral LPFC had a significantly different proportion of neurons tuned to these features (Chi-Square, χ = 12.20, p < 0.05 in monkey B, Chi-Square, χ = 30.36, p < 0.001 in monkey T).

**Figure 2.**
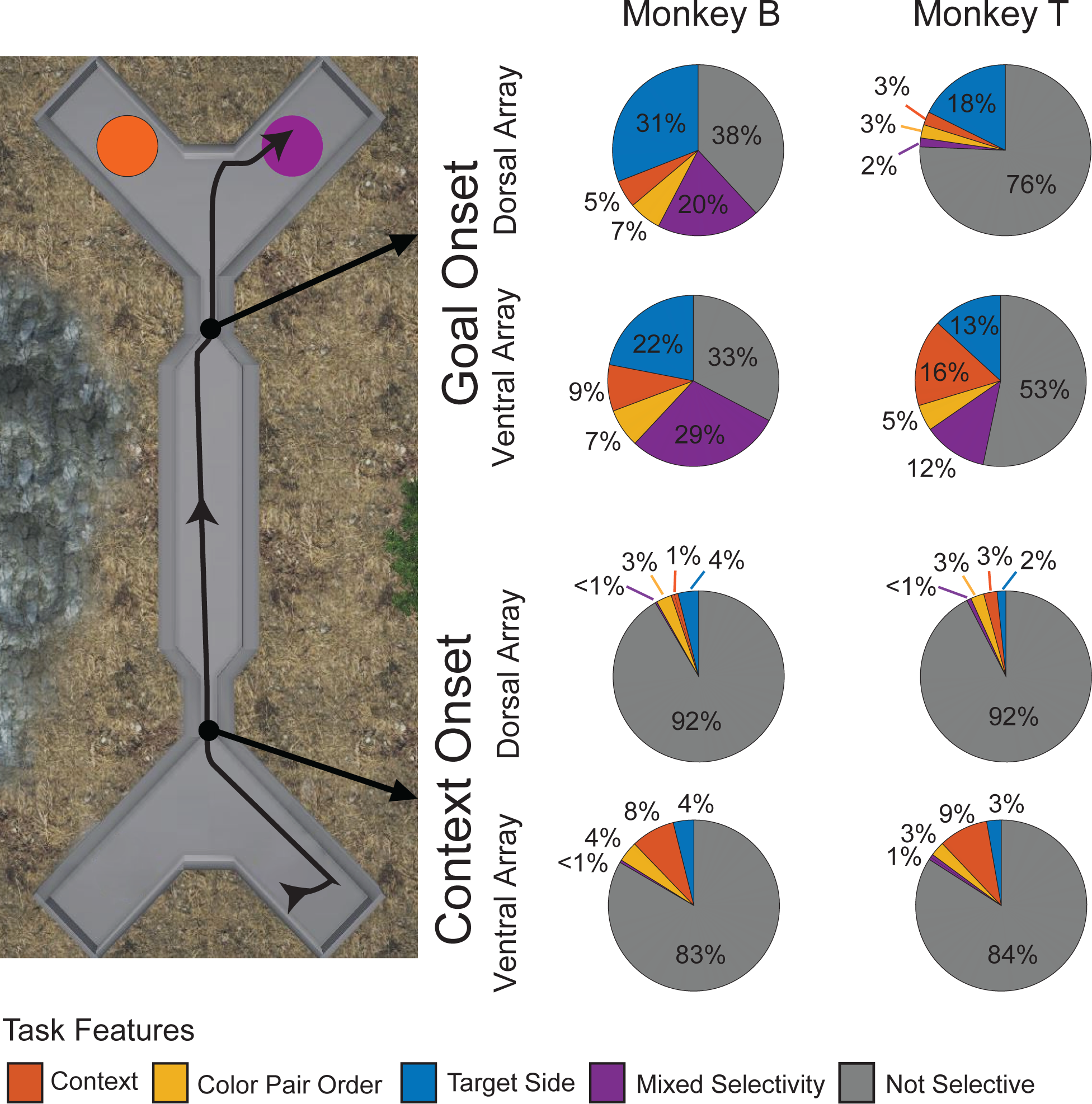
LPFC neurons are tuned to task features during virtual navigation. Proportion of neurons tuned to task features during the context onset and goal onset epochs, using a multivariate linear regression over the mean normalized firing rate of each neuron over 1200ms (-200ms to 1000ms) around each epoch for both monkeys. An alpha of 0.05 was used to determine neuron tuning.

### LPFC neurons temporally mix task features during virtual navigation

During the onset of the goal, the monkeys must associate the context with the appropriate color and navigate towards it. The naturalistic features of this task (e.g., gaze was unconstrained, and the animal initiated and conducted navigation at will using a joystick) allowed the monkeys to freely complete the task and navigate towards their goal, with no additional behavioural constraints. This provided us with the opportunity to explore neural dynamics in the LPFC which may take place while performing real-world context-target associations. To this end, we investigated neuron tuning around the goal onset epoch by performing multivariate linear regressions over a 400ms window running in steps of 20ms. Each unit was identified as being tuned to a feature if it remained significantly tuned over five consecutive steps.

In both monkeys, a significant proportion of neurons were tuned for task features (**Figure 3a**). Neurons in the ventral LPFC acquired tuning to task features in the same temporal pattern: first context, then CPO, and finally target side (**Figure 3b**). A significant proportion of neurons were tuned to the context at all times investigated, with a mean latency (± standard error) of 130.1 ± 27.4 ms in monkey B and 118.2 ± 31.7 ms in monkey T. Individual neurons then demonstrated CPO tuning significantly later relative to context (296.2 ± 21.4 ms, p<0.001 in monkey B, 290.3 ± 25.2 ms, p<0.01 in monkey T, Wilcoxon rank sum). Neurons finally acquired tuning to the target side (493.3 ± 19.3 ms, p<0.001 in monkey B, 407.5 ± 26.2 ms, p<0.05 in monkey T, Wilcoxon rank sum). Neurons in the dorsal LPFC had different tuning profiles in the two monkeys. Both animals had neurons that were tuned to target side (450.7 ± 15.9 ms in monkey B, 528.7 ± 29.6 ms in monkey T). However, only monkey B had neurons in the dorsal LPFC tuned to context (214.2 ± 22.5 ms) followed by CPO (325.7 ± 15.4 ms, p<0.001, Wilcoxon rank sum). Many of the same neurons were tuned to different features over the course of the trial, demonstrating time multiplexing of their mixed selectivity (**Figure 3c**). We performed the same analysis with different alpha thresholds (0.01 and 0.001). Although this decreased the proportion of tuned units, our overall findings did not change (**Figure S1a**).

**Figure 3.**
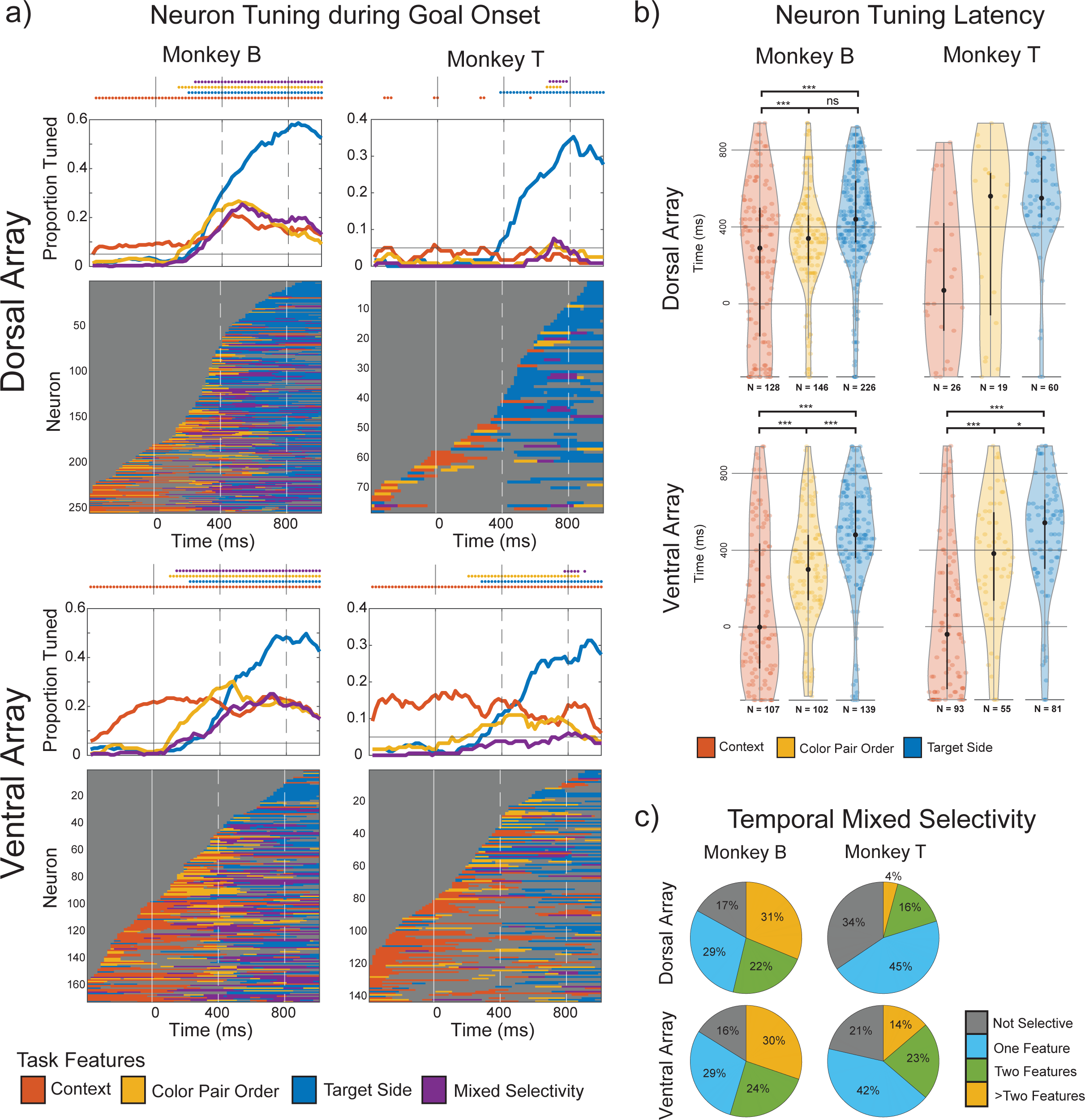
LPFC neurons temporally mix task features during virtual navigation. a) Neuron tuning to task features during goal onset over 400ms windows overlapped by 20ms, performed separately across the ventral and dorsal PFC for each monkey. Top plots demonstrate the total number of neurons tuned to task features, with the points above indicating a proportion of units tuned above 0.05 (see Figure S1 for results with an alpha of 0.01 and 0.001). Bottom plots represent each unit’s tuning over time, sorted from bottom to top by onset of tuning to any feature. b) Latencies of all neurons tuned to task features in the ventral and dorsal PFC across both monkeys, with latencies compared using a Wilcoxon rank test, ns p>0.05, * p<0.05, *** p<0.001. c) Proportion of neurons tuned to different features over the 1200ms (-200ms to 1000ms) time-window around goal onset.

To mimic natural behaviour, we did not constrain eye position in our virtual reality task. Thus, eye position or the allocation of attention (e.g., overt attention) associated to it could have had an influenced in the neural responses. To investigate this issue, we performed a two-step multivariate linear regression (see methods). We first calculated the mean x and y eye positions in the same 400ms windows used to analyze the neural data and fit the eye position to the normalized firing rates (FRs, Equation 2 in methods). The residual FR error (FR∈) was then used in a second multivariate linear regression model that used task-related features as independent variables (Equation 3 in methods). The total proportion of neurons tuned to task features following this analysis are presented in **Figure S1b**. Both monkeys had a bias to fixate on their chosen side, and monkey B had a bias to preferentially fixate on one of the two colored objects presented in some sessions (**Figure S2**). Therefore, removing eye-position related information resulted in fewer neurons being tuned to target side in both monkeys, and fewer neurons tuned for CPO neurons in monkey B due to his color preference bias. Despite this, we observed the same temporal pattern previously described in the ventral LPFC in both monkeys. Remarkably, unlike in the ventral LPFC, monkey B’s dorsal LPFC had no significant CPO tuned neurons until 380ms after the goal onset. This further suggests that unlike in the ventral LPFC, neurons in monkey B’s dorsal LPFC do not encode CPO prior to the target side or the allocation of attention/gaze to that side.

### Decoding task features of ventral and dorsal LFPC neuronal ensembles across individual sessions

We found that individual neurons are dynamically tuned to task features. However, single neuron analysis may not adequately capture the informational capacity of a population of simultaneously recorded units (Tremblay et al. 2015). We therefore used a linear support vector machine (SVM) to classify task-features from neuronal ensemble activity. We performed this analysis on each session separately, using sessions with at least 15 correct trials for each of the four conditions. We used firing rates integrated over the same 400ms windows in steps of 20ms around goal onset.

We trained and tested a linear SVM on normalized thresholded multiunit activity on each channel of the dorsal and ventral LPFC arrays (see methods). The mean (±2 standard error, SE) decoding performance across 12 sessions in Monkey B and 8 sessions in Monkey T is presented in **Figure 4a**. Each point is marked when the when the lower bound (mean – 2SE) exceeds chance performance. Task features could be reliably decoded following the same temporal pattern as observed with single neuron tuning. This suggests ensembles in the ventral LPFC represented context information throughout the task in both monkeys (orange traces in **Figure 4a**). Additionally, CPO could be decoded from the ventral LPFC after the goal onset, followed by the target side. Only target side could be decoded from the dorsal LPFC in monkey T (right upper panel). However, context and CPO information could be decoded from the dorsal LPFC in monkey B (left upper panel). These results were observed across individual sessions in monkeys B and T (**Figure S3**). Permutation testing with shuffled labels was used to obtain a 95% confidence interval for individual sessions (see methods). We repeated the same ensemble analyses only including well isolated single neurons. There was no significant difference (Wilcoxon rank sum test) in decoding performance when training the classifier on multiunit activity and manually sorted spikes for both monkeys (**Figure S4**).

**Figure 4.**
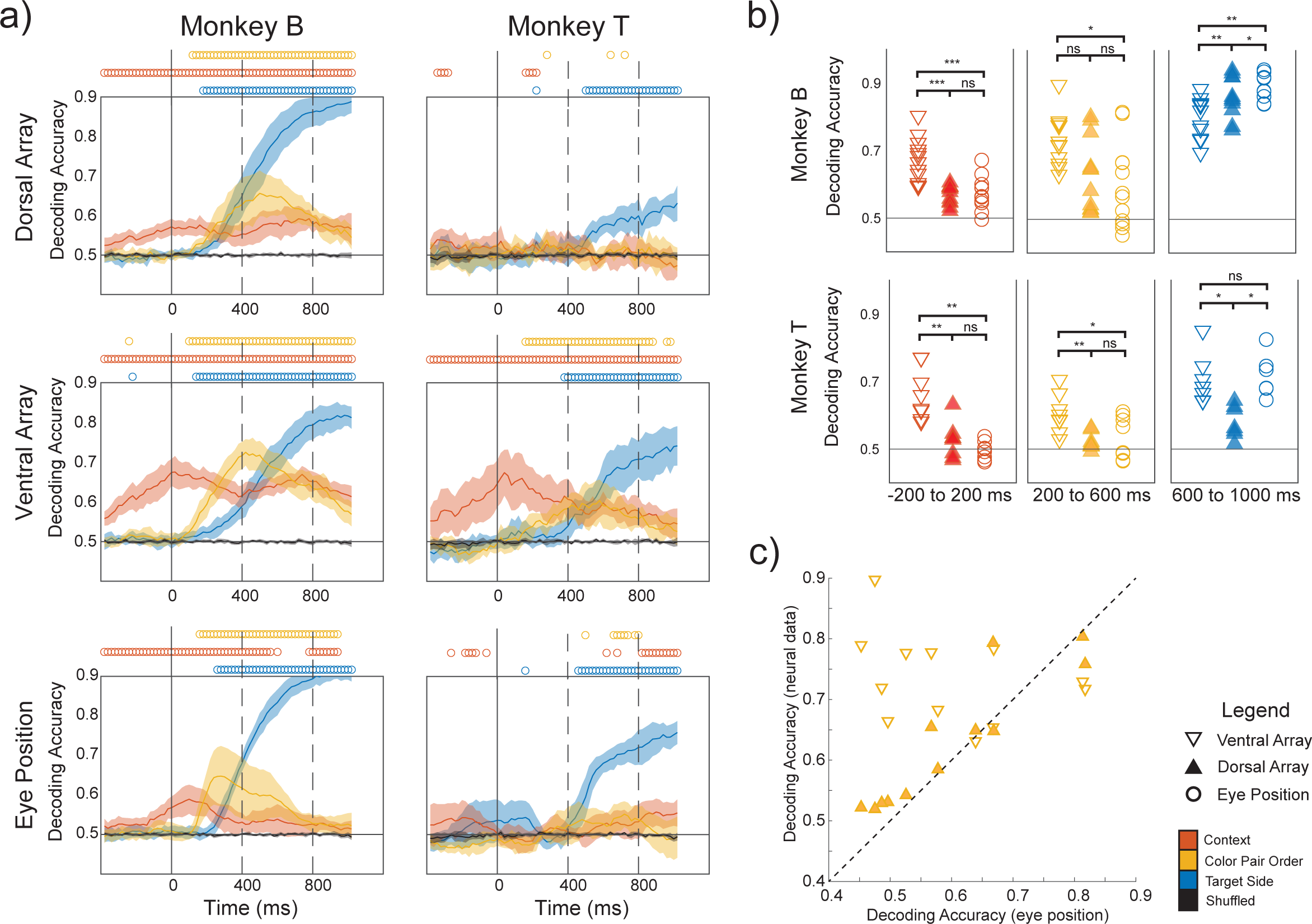
Decoding task features of ventral and dorsal LFPC neuronal ensembles across individual sessions. a) Mean decoding accuracies of task features across 12 sessions (monkey B) and 8 sessions (monkey T) over 400ms windows overlapped by 20ms. This was performed separately for neural activity in the dorsal and ventral LPFC (mean and normalized multiunit activity), and mean eye position (mean x and y positions) over the same time epochs. Plots show mean (±2SE) decoding accuracy across sessions, with a circle above when the lower bound (mean – 2SE) exceeds chance performance. b) Comparing decoding accuracy across the ventral LPFC, dorsal LPFC and eye position using a Wilcoxon sign rank test, ns p>0.05, * p<0.05, ** p<0.01, *** p<0.001. c) Correlating decoding accuracy of CPO from eye position and neural data 400ms (200ms to 600ms) after goal onset using Spearman’s rank correlation. Decoding accuracy of CPO from eye position was highly correlated with accuracy from the dorsal LPFC (r(12) = 0.918, p < 0.001), but not from the ventral LPFC (r(12) = -0.329, p = 0.30).

We then compared decoding accuracies of task features from the dorsal LPFC, ventral LPFC and eye position using a Wilcoxon sign rank test at three 400ms intervals centered at 0ms, 400ms and 800ms after goal onset (**Figures 4b and S5**). Context information could be decoded more accurately from the ventral LPFC compared to eye position (p<0.001 in monkey B, and p<0.01 in monkey T, Wilcoxon sign rank), and the dorsal LPFC (p<0.001 in monkey B, and p<0.01 in monkey T, Wilcoxon sign rank). CPO could also be more accurately decoded from the ventral LPFC compared to eye position at 400ms (p<0.05 in monkey B, and p<0.05 in monkey T, Wilcoxon sign rank), and the dorsal LPFC (p=0.05 in monkey B, and p<0.01 in monkey T, Wilcoxon sign rank). Given monkey B’s bias to fixate on one colour in some sessions (**Figure S2**), we performed a Spearman’s rank correlation between decoding accuracy from eye position compared to neural activity (**Figure 4c**). CPO information was independent of eye position only in the ventral LPFC at 400ms, consistent with our single neuron findings (**Figure S1b**).

### Task features are fluidly represented in the ventral LPFC

Our population level analysis shows neural ensembles represent task-features following the flow of information during task trials or the cognitive operations the animal needs to perform the task. Individual neurons also fluidly represented task features, with different neurons acquiring or modulating their tuning throughout a trial (**Figure 3**). These temporal multiplexing of features would predict that the neural codes across the different trial periods are dynamic and likely do not generalize from one period to another. To investigate this issue, we trained a linear SVM to classify task-features at each 400ms window and tested this at the other different time windows (**Figures 5**, **S6** and **S7**).

**Figure 5.**
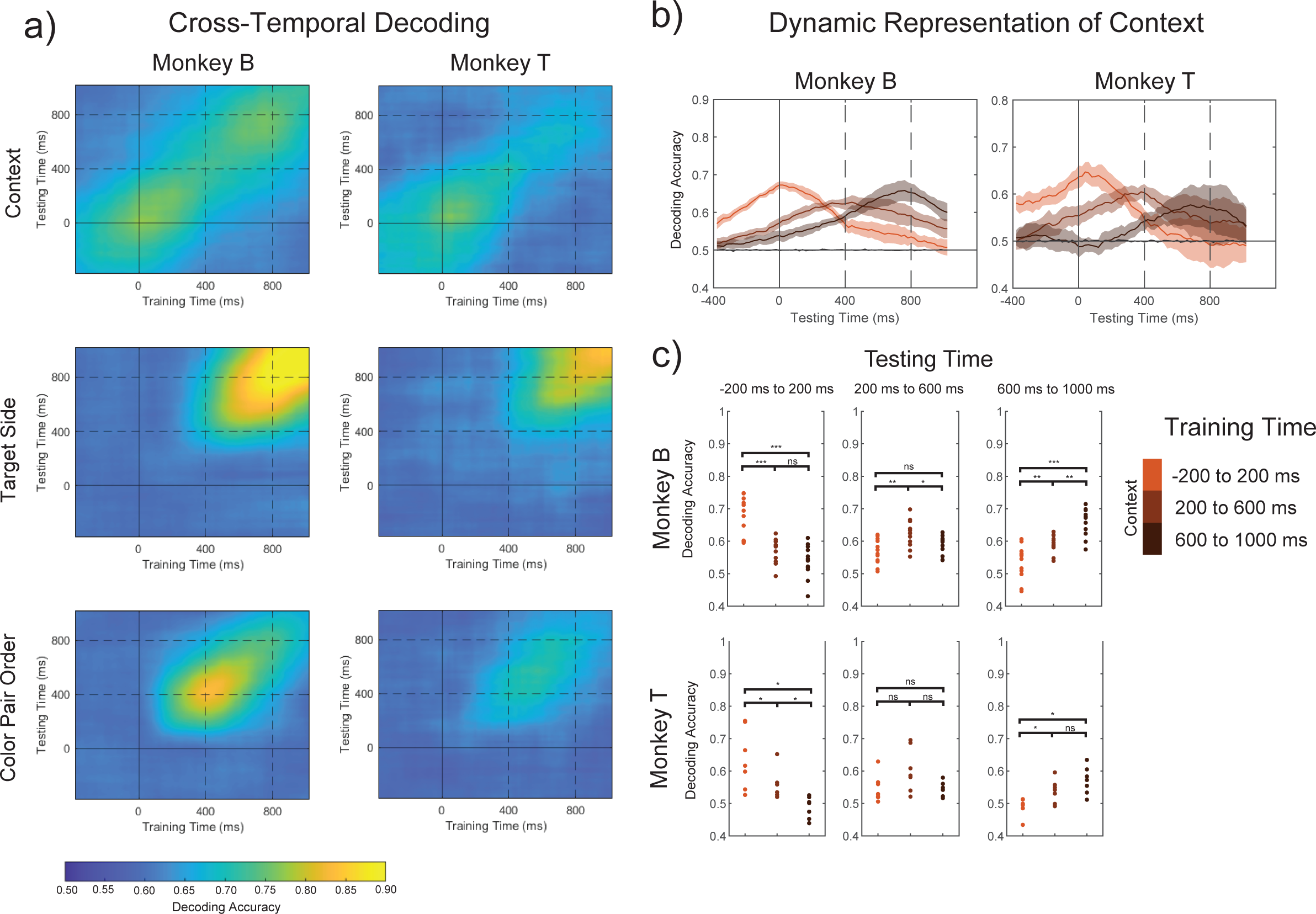
Task features are fluidly represented by neural ensembles in the ventral LPFC. a) Cross-temporal decoding in the ventral LPFC, training a linear SVM on task features across 400ms windows overlapped by 20ms, and testing each model on 400ms windows overlapped by 20ms across the whole epoch. Heatmaps show mean decoding accuracies across 8 sessions in monkey B and 8 sessions in monkey T. b) Mean (±2SE) decoding accuracy of the task’s context across time, when trained at 0ms, 400ms and 800ms. c) Comparing decoding accuracies of models trained at 0ms, 400ms and 800ms when testing on different time windows. Decoding accuracies were compared using a Wilcoxon sign rank test, ns p>0.05, * p<0.05, ** p<0.01, *** p<0.001.

Context information was represented with a dynamic neural code (**Figure 5a** upper panels and **Figure 5b**). Context could be more accurately decoded at 0ms from goal onset when the classifier was trained at that time compared to later in the epoch at 400ms and 800ms in both monkeys (p<0.001 in monkey B, p<0.05 in monkey T, Wilcoxon sign rank) (**Figure 5c**). Conversely, context was more accurately decoded at 800ms from a classifier trained at that time compared to one trained at 0ms (p<0.001 in monkey B, p<0.05 in monkey T, Wilcoxon sign rank) (**Figure 5c**). Target side information was represented by a more persistent code (**Figure 5a** middle panels), with no difference in decoding accuracy at 400ms when trained at that time or at 800ms in both monkeys (**Figure S6** and **S7**). Finally, CPO demonstrated a partially dynamic neural code (**Figure 5c** middle column). Training a linear decoder to classify this feature at 400ms demonstrated increased decoding accuracy at that time compared to a classifier trained at 800ms (p<0.01 in monkey B, p<0.05 in monkey T, Wilcoxon sign rank); however, testing at 800ms demonstrated the same accuracy when trained at 400ms and 800ms (p>0.05 in both monkeys, Wilcoxon sign rank). This suggests that there was a greater representation of CPO at 400ms, but there was no additional CPO information present at 800ms. To rule out a bias in eye position influencing this feature in monkey B, we only used sessions (n=8) where he had no significant bias to fixate on one colour. Although this limits the bias of eye position, it likely does not completely remove it and may explain some of the cross-temporal decoding differences observed in both the dorsal and ventral LPFC (**Figure S6**).

To corroborate whether this dynamic code was linked to the dynamic nature of individual neuron tuning, we compared the weights of our linear SVM for context to the regression coefficients (betas) from our single neuron multivariate regressions previously described (**Figure 3**). As anticipated, there was a significant correlation between the regression coefficients of individual neurons tuned to the task-features and their weights determined by our linear decoder (**Figure 6).** This suggests our linear decoder uses the same neurons that are tuned at each time window. This stresses the fluid nature of the task but also a degree of serial information processing by single neurons and ensembles.

**Figure 6.**
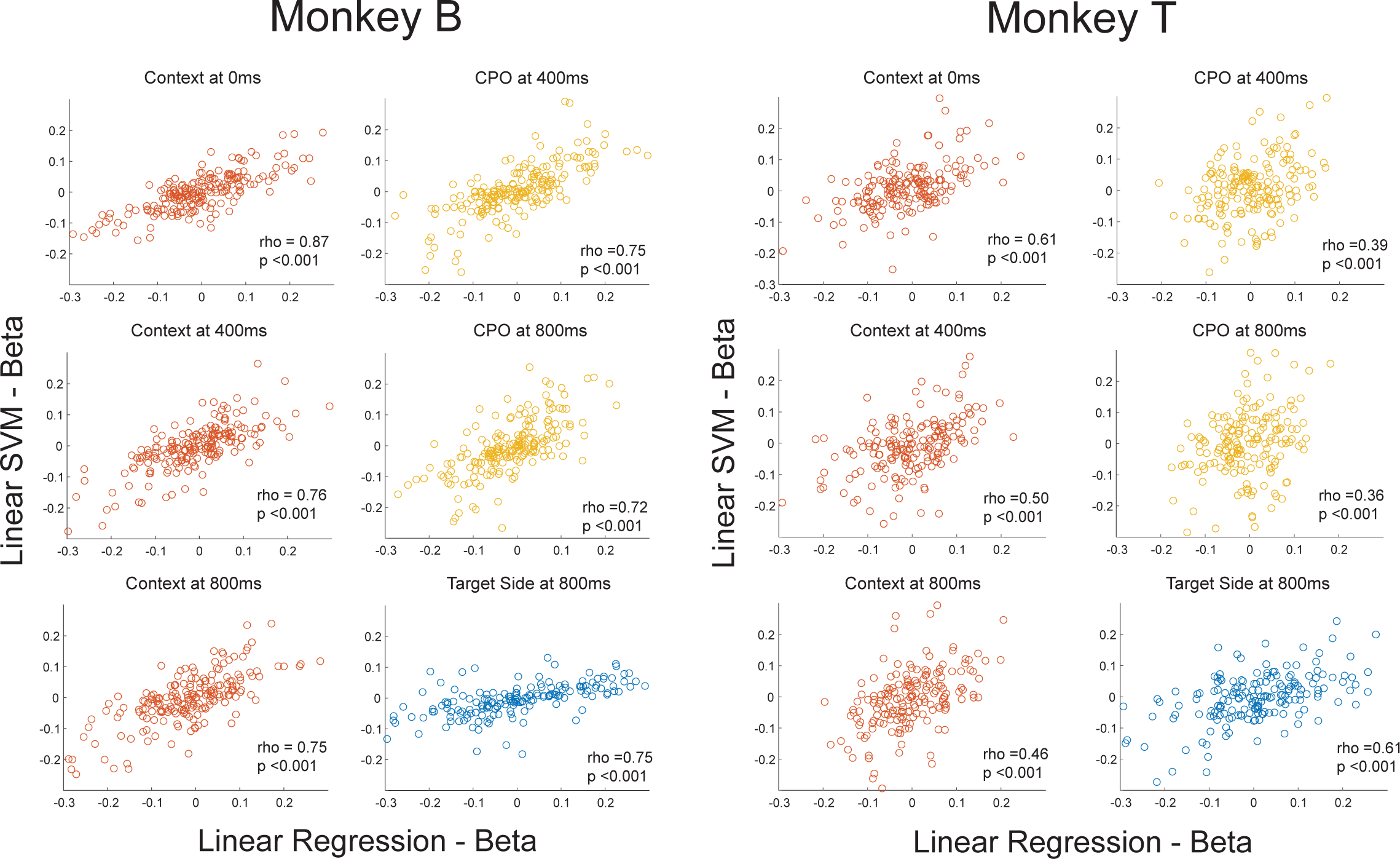
Temporally mixed single neurons contribute to fluid neural ensembles. Correlating each sorted unit’s beta coefficient for context, CPO and side obtained using a multivariate linear regression as previously described (Figure 3), to each unit’s beta coefficient as assigned by a linear SVM classifying context, CPO and target side at different times. Correlation was assessed using a Spearman rank correlation, with rho and p-values presented.

## Discussion

We recorded single neuron responses in the dorsal and ventral LPFC of macaques while the animals navigated a virtual environment and performed an associative memory task with variable spatiotemporal structure. Our main results were 1) single neurons in the LPFC show mixed selectivity for task relevant features, 2) single neurons and ensembles time-multiplexed their selectivity for the different elements of the task depending on the time-relevance of that element to solve the task, and 3) while neurons in the ventral LPFC encoded spatial and non-spatial features, neurons in the dorsal LPFC encoded spatial information.

### Neural dynamics in a naturalistic setting

Few studies have examined single neuron tuning in the primate LPFC in naturalistic settings. During most experiments, eye position is strictly controlled and stimuli are presented on a stationary background. In these studies animals do not have the ability to change their view or position in the environment and therefore cannot control the spatiotemporal dynamic of task elements. These restrictions have served as important controls limiting experimental confounds, but also limit the ability to generalize findings to many real-world scenarios. We reasoned the LPFC must perform its function in complex environments in the presence of gaze movements and changing scenery. One study (Tremblay et al., 2023) trained monkeys to perform a simple associative memory task without limiting the animals’ head and eye movements. They showed that coding of relevant features in the LPFC was robust and not degraded by unstructured movements. Although this work is an important step towards understanding neural representations in naturalistic settings, it did not explore neuron tuning in a complex environment such as the one in our task where features changed as the animal voluntarily moved through the environment. This capacity to execute actions that cause a change in the virtual world is referred to as a feeling of agency (Bódi, 2022). Other studies have used virtual environments as a surrogate for real-world naturalistic settings and reported tuning for spatial working memory in LPFC and for stimulus features in the hippocampus (Corrigan et al., 2022; Gulli et al., 2020; Johnston et al., 2023; Roussy et al., 2021; Wirth et al., 2017). This factor should not be underestimated since here animals do not act as passive observers but as actors, triggering task events.

Unconstrained naturalistic tasks in which subjects have agency may provide insights into the neural dynamics of the LPFC during real-world conditions. We showed that single neurons in the LFPC were serially tuned to relevant task features as they ‘appear’ in a naturalistic associative memory task: first representing context, followed by the CPO, and finally target location. Interestingly, neurons largely acquired tuning to the context shortly before the goal was triggered (**Figure 3**), despite this information being previously available, suggesting context was only represented in the LPFC when becoming immediately relevant. Furthermore, these neural representations were fluid over time, both at the single neuron (**Figure 3**) and ensemble level (**Figure 5**). Single neurons in the LPFC have been observed to non-linearly mix task-relevant features, allowing for flexible high-dimensional representations that can be read out from downstream cortical areas (Asaad et al., 1998; Dang et al., 2021; Fusi et al., 2016; Rigotti et al., 2013). Our results support these findings and further show that LPFC neurons multiplex features in time as they become task relevant. This property would allow a single component of mixed selectivity to be represented over trial periods or entire trials. Possible advantages of time multiplexing are utilizing the same neurons to represent individual features as well as combinations of those features united by rules (Buckley et al., 2009; Mendoza-Halliday & Martinez-Trujillo, 2017; Tanji & Hoshi, 2008; Wallis et al., 2001), and a fluid representation of a spatiotemporal episode within the same population of neurons in the LPFC (Roussy et al., 2021). The LPFC has been proposed to serve as a buffer for episodic working memory (Baddeley, 2000). The time multiplexing in this region may be critical for the spatiotemporal binding of stimulus features required during the perception of a memory episode.

### Anatomical and functional subdivision of LPFC

The neural dynamics previously discussed was mostly found in in the ventral LPFC in both monkeys tested, suggesting a functional division withing this region. The ventral and dorsal LPFC had different tuning profiles while the monkeys completed the task, with more neurons in the ventral LPFC being tuned to visual and visuospatial features. The ordered tuning of context, target location and target side was observed only in the ventral LPFC. These results suggest the ventral LPFC is more likely to represent visual features as compared to the dorsal LPFC. These results are consistent with connectivity studies that show there is dorsal and ventral mapping of the LPFC to corresponding dorsal and ventral high-level visual areas (Barbas, 2015; Barbas & Pandya, 1989; Petrides, 2005b; Xu et al., 2022).

Other single neuron studies have demonstrated more mixed results regarding a functional organization to the LPFC (Asaad et al., 1998; Kadohisa et al., 2015; Meyer et al., 2011; O’Reilly, 2010; Riley et al., 2016; Siegel et al., 2015; Tang et al., 2021; Wallis et al., 2001). In contrast, we observe a relatively robust difference between single neuron tuning and ensemble dynamics between the ventral and dorsal LFPC. The increased cognitive demands of interpreting a complex scene and freely navigating through it may exacerbate underlying regional specializations in function. This is the first study to examine anatomical differences in LPFC function at the single neuron and population level during a virtual navigation task in which the subjects have agency. Importantly, our study demonstrates it is possible to use a virtual task to explore information processing in monkey LPFC neurons and offers novel insights into the dynamic nature of their mixed selectivity.

Our results have implications for models of LPFC function and interactions with the rest of the brain. Interestingly, one recent study has shown that stimulation of the LPFC evokes selective activation of specific regions in associative areas of the neocortex (Xu et al., 2022). The connectivity of LPFC with early sensory areas was practically non-existent suggesting the LPFC mainly receives integrated information from association areas. Because the mixing of features is not solely a feature of the LPFC but may start in association cortices, the information that LPFC neurons integrate could be highly filtered and features may be already mixed. The latter may facilitate the time multiplexing of relevant features observed in our study as well as how feedback signals from the LPFC may reach association areas. Thus, the reported role of the LPFC on several components of cognitive control such as attention (Backen et al., 2018; Bichot et al., 2015; Duong et al., 2019; Lennert & Martinez-Trujillo, 2011), working memory (Roussy et al., 2021), perception (Mendoza-Halliday et al., 2018), rule coding (Buckley et al., 2009; Corrigan et al., 2022; Kaping et al., 2011), planning (Tanji et al., 2007) (see (Miller, 2000) for a review) could be understood in neurophysiological terms as containing populations of neurons that can serve as spatiotemporal integrators of information held in the focus of perceptual awareness. This may enable the high level of mental abstraction observed in anthropoid primates (Passingham & Wise, 2012) with larger prefrontal cortices.

## Methods

### Experimental Animals

Two male rhesus macaques (*Macaca mulatta*; 7 and 14 years old, 7kg and 12kg respectively) were used in these experiments. The monkeys were trained to perform a virtual reality associative memory task and received a juice reward as a form of positive reinforcement for each session. All animal procedures were compliant with the Canadian Council on Animal Care guidelines and approved by the Western University Animal Care Committee.

### Electrophysiological recordings

Lateral prefrontal cortical (LPFC) recordings were acquired using two 96-channel Utah arrays for each animal (Blackrock Microsystems). Each animal received a 3-Tesla T1-weighted MRI which was used for surgical navigation using Brainsight (Rogue Research Inc.). The Utah arrays were positioned just anterior to the arcuate sulcus, with one placed dorsal to the principal sulcus (areas 46d/9d) and one placed ventral to the principal sulcus (areas 46v/9v). Each shank was 1.5mm and was therefore likely in layers II/III. Intraoperative images of the arrays’ positions for each animal are presented in **Figure 1a**, with the corresponding regions highlighted in the MRI scans. Signals were obtained at 30kHz using a Cerebus Neural Signal Processor (Blackrock Microsystems) and saved for offline sorting. Extracted spikes were semi-automatically sorted using Plexon Offline Sorted (Plexon Inc.), and then manually sorted by two raters (BWC and MA).

### Experimental setup

The animals completed the task while seated in front of a computer monitor (27” ASUS VG278H monitor, 1024x768 pixel resolution at a 75 Hz refresh rate) and used a two-axis joystick to freely navigate through the virtual environment. They were placed in a radiofrequency shielded dark room, with cables entering the room through a small aperture. The monkeys’ head position was fixed during the task, and eye position was recorded using video-oculography at a sampling rate of 500Hz (EyeLink 1000, SR Research). Custom software was used to simultaneously control and record the stimulus presentation, behavioral response, eye position and reward dispense **(Figure 1a**).

### Behavioral tasks

The learning task was completed in an X-shaped maze as previously described (Corrigan et al., 2022; Doucet et al., 2016; Gulli et al., 2020). In each trial, animals start at one end of the X-maze and navigate towards the corridor using a joystick. Once they enter the corridor, the walls change their texture to either a wood or steel texture. As they continue navigating towards the end of the corridor, two different colored discs appear at each end of the X-maze. One wall texture (the context) is associated with one color (e.g., steel means the target is green, and wood means the target is red) indicating the monkeys to navigate to the corresponding disc to obtain a juice reward. The monkey then turns around to navigate back towards the other end of the maze initiating another trial. **Figure 1c** demonstrates the trajectory of an example trial.

Each day, monkeys completed a set of trials with a fixed context-color combination, which associated the steel context with the orange target, and the wood context with the purple target. Following this fixed combination, two different colors are pseudo-randomly chosen to be associated with each context. Therefore, the monkeys must learn a new association for each session. For a given trial, the colors can appear in one of two possible configurations, referred to as color pair order (CPO). **Figure 1d** demonstrates the different possible task elements for each session: context, CPO and target side. For a given trial, the context and CPO are chosen randomly.

### Behavioral analysis

For each session, learning was estimated using a state-space analysis based on all completed trials as previously described (Ravassard et al., 2013). A learning state with a 95% confidence interval is estimated across all completed trials, as illustrated in **Figure 1e**. Learning of the association is defined as the 2.5^th^ percentile of the learning state’s confidence interval exceeding 50%.

### Single neuron tuning

Single neuron tuning over task epochs was computed by calculating the mean firing rate (FR) over 1200ms, from -200ms to 1000ms around the trial’s epoch. Each unit’s mean FR was z-scored across all trials. Given a limited number of incorrect trials, only correct trials were used. We fit each neuron’s normalized FR to each feature using a multivariate linear regression (*equation 1*), with significance to each feature determined by setting an alpha of 0.05. The total proportions of neurons tuned to task features were compared across the dorsal and ventral LPFC using a chi-square test.

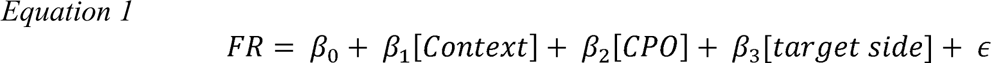

In the model *context is the wall color (2 values: wood or steel), CPO is color pair order,* the color combination order *(2 values: color1 left and color2 right and vice versa)*, and target side is the location of the target in egocentric space (left or right). To investigate neuron tuning to features over the task epoch, we calculated for each unit the mean FR over 400ms windows with 20ms overlap. This was calculated from -400ms (-200ms to -600ms) to 800ms (600ms to 1000ms) around the goal onset. Each unit’s mean firing (FR) rate z-scored across all trials and tuning to task features was determine using a multivariate linear regression (Equation 1). We perform this analysis with an alpha threshold of 0.05, 0.01 and 0.001. A given unit was defined as being tuned to a feature if it remained tuned for at least five consecutive 20ms sliding windows (i.e. across 100ms). The latency for a given unit’s tuning was determined as the first time-window it was significantly tuned to a given feature. The total number of tuned units for each time-window were summed for each feature, and a significant proportion of neurons was determined if the total proportion exceeded the alpha threshold set. Individual neuron tuning latencies were compared for different task features using a Wilcoxon rank sum test.

Eye position was unrestrained. To control for eye position, we performed a two-step multivariate linear regression (Equations 2 and 3). We first calculate the mean x and y eye position in the same 400ms windows we used to analyze the neural data. We fit each unit’s normalized FR to the mean x and y eye position, and their interaction to account for neural activity specific to a particular position on the screen (Equation 2). We then utilize the residual FR data, representing the neural activity not accounted for by eye position, to fit a second multivariate linear regression with task-dependent features (equation 3).

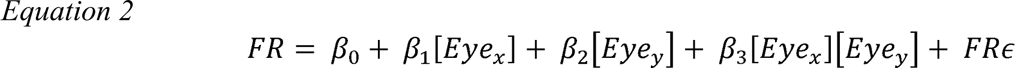

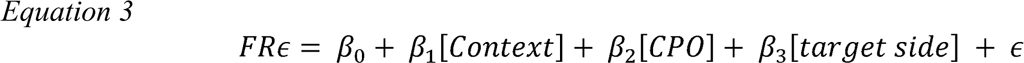

### Decoding from Population Neural Activity

We used a linear support vector machine (SVM) to decode task-dependent features from neuronal population activity in single trials in 400ms windows. We initially used only correct trials, with four possible conditions. We utilized the fitcsvm function (MATLAB 2020a) using a five-fold cross validation on normalized FRs. Sessions were analyzed separately, and only sessions with at least 15 trials per condition were used. For each session, 15 trials for each condition were randomly chosen and stratified into five folds with one-fold used for testing. This procedure was repeated 100 times, with 500 trained classifiers. In individual sessions, significance was determined by permutation testing, with the previous procedure performed on shuffled trials. Trials were shuffled after the k-fold stratification, and this procedure was repeated 100 times, with 500 classifiers. In individual sessions, significance was determined by a mean accuracy exceeding the 97.5^th^ percentile chance accuracy. We performed the above analysis on normalized FRs from sorted neurons and normalized multiunit activity across each channel.

To control for possible biases in eye position that may affect neural activity we sought to quantify the classification accuracy of task-features from eye position. We calculated the mean x and y eye position across the same 400ms windows as previously described. A linear SVM was used to classify task-features from x and y eye position on the same randomly selected and stratified trials used for the neural data. This was repeated 100 times, generating 500 classifiers. Permutation testing as described above was used for significance testing in individual sessions. The mean decoding accuracies (with standard errors) of task features were calculated across sessions. Decoding accuracies were compared across sessions using the dorsal and ventral LPFC’s neural data and eye position at three 400ms windows (centered at 0ms 400ms and 800ms) using Wilcoxon sign rank tests.

### Cross-temporal Decoding Analysis

Cross-temporal decoding was performed to investigate whether the neural code underlying task features was persistent or dynamic across the goal onset epoch. We calculated the mean FR across 400ms windows overlapped by 20ms, and randomly selected trials which were stratified in five k-folds. This was performed from -400ms to 1000ms around the goal onset, with 70 separate time-windows. A linear SVM was used to generate a model on 4 k-folds at each time-window and tested on the same remaining k-fold across all 400ms time-windows (70 time windows). This procedure was then repeated 100 times, resulting in 35 000 separate models (70 x 5 x 100). The mean decoding accuracy across sessions (with the standard error) was calculated, and significance was determined when the bottom 2.5^th^ percentile decoding accuracy exceeded chance performance. Cross-temporal decoding analysis was statistically compared across models trained at three time windows (centered at 0ms, 400ms and 800ms) around goal onset using a Wilcoxon sign rank test.

## Supplementary Figure Captions

**Figure S1. Tuning of LPFC neurons using different alpha thresholds and correcting for eye position.** a) Proportion of neurons tuned to task features during goal onset over 400ms windows overlapped by 20ms, using different alpha thresholds (0.01 and 0.001). The points above indicating the proportion of units tuned above the expected number tuned by chance based on the alpha set. b) Neuron tuning to task features during goal onset over 400ms windows overlapped by 20ms, after removing eye position related information using a multi-step multivariate linear regression. Total proportion of neurons tuned plotted above, with points above indicating a proportion of units tuned above the alpha set (0.05).

**Figure S2. Examining mean eye position in select sessions across both monkeys.** a) Mean horizontal eye position over running 400ms windows overlapped by 20ms across task conditions with a 95% confidence interval shaded, for individual select sessions. Sessions B3 and B5 were selected for monkey B as they sessions had the most and least eye-position bias for task features respectively, and session T5 is shown as this session had the most trials in monkey T. b) Decoding accuracies of task features using mean x and y eye position over 400ms in individual sessions shown above, with the shaded gray area demonstrating the 95% confidence interval obtained by shuffling the labels. Chosen side can be decoded above chance level in all sessions as the monkeys fixated on the target side. Session B3 demonstrates robust decoding of Color Pair Order, as monkey B had a preference to fixate on the cyan color (and therefore was looking in a different direction depending on the color order). However, monkey B did not have a color preference in session B5 and therefore Color Pair Order could not be decoded above as robustly in this session. Monkey T did not demonstrate a similar bias for visual task features, and in fact did not reliably fixate on the target side in all trials.

**Figure S3. Decoding task features from individual sessions.** Mean decoding accuracy of task features in individual sessions with at least 30 trials in each condition, with the shaded gray area representing the 95% confidence interval obtained with permutation testing.

**Figure S4. Decoding task features using manually sorted single units and multiunit activity.** Mean decoding accuracy (±2SE) of task features using normalized sorted single neuron spikes (left) and multi-unit activity across each channel (right), over 400ms time windows overlapped by 20ms. Decoding accuracies were statistically compared at 0ms, 400ms and 800ms using a Wilcoxon rank sum test, with no statistical difference observed between manually sorted single units and multiunit activity.

**Figure S5. Decoding accuracies of task features between neural data from the dorsal and ventral LPFC and eye position.** Decoding accuracies of task features by the dorsal PFC, ventral PFC and eye position compared at three time windows (centered at 0ms, 400ms and 800ms). Decoding accuracies across sessions were compared using a Wilcoxon sign rank test, ns p>0.05, * p<0.05, ** p<0.01, *** p<0.001.

**Figure S6. Cross-temporal decoding from the dorsal and ventral LPFC performed in Monkey B.** a) Heatmap representing cross-temporal decoding accuracies, with a linear SVM trained in 400ms intervals overlapped by 20ms, and tested at all other 400ms windows. b) Mean decoding accuracy over testing time when training at 0ms, 400ms and 800ms. c. Comparing decoding accuracies of models trained at 0ms, 400ms and 800ms when testing on different time windows, using a Wilcoxon sign rank test, ns p>0.05, * p<0.05, ** p<0.01, *** p<0.001.

**Figure S7. Cross-temporal decoding from the dorsal and ventral LPFC performed in Monkey T.** Cross-temporal decoding in monkey T a. Heatmap representing cross-temporal decoding accuracies, with a linear SVM trained in 400ms intervals overlapped by 20ms, and tested at all other 400ms windows. b. Mean decoding accuracy over testing time when training at 0ms, 400ms and 800ms. c. Comparing decoding accuracies of models trained at 0ms, 400ms and 800ms when testing on different time windows, using a Wilcoxon sign rank test, ns p>0.05, * p<0.05, ** p<0.01, *** p<0.001.

## Notes

### Competing Interest Statement

The authors have declared no competing interest.

