## Supplemental Figures for "Neural Ensembles in the Lateral Prefrontal Cortex Temporally Multiplex Task Features During Virtual Navigation"

### Supplementary Figure 1

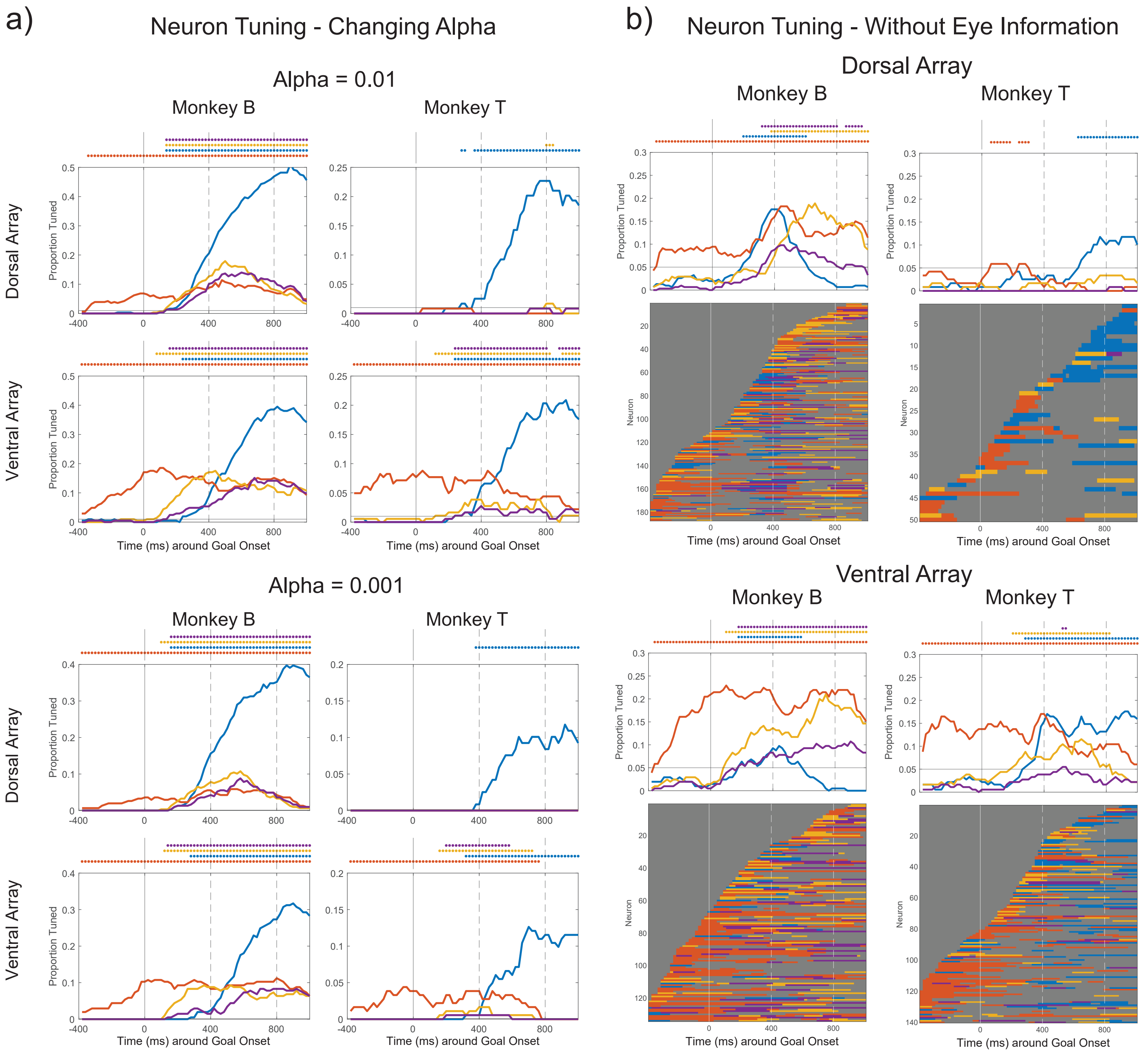

### Supplementary Figure 2

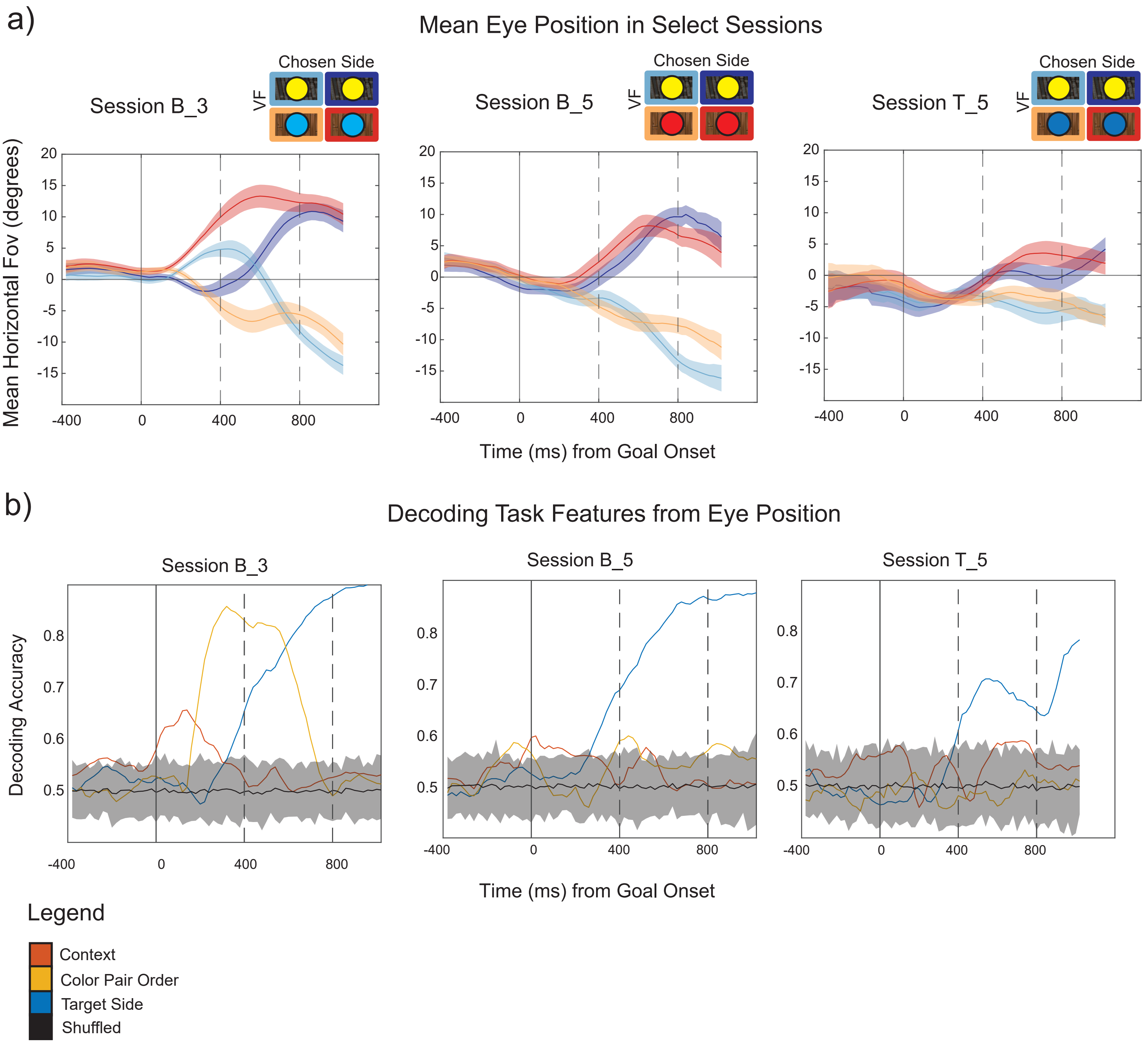

### Supplementary Figure 3

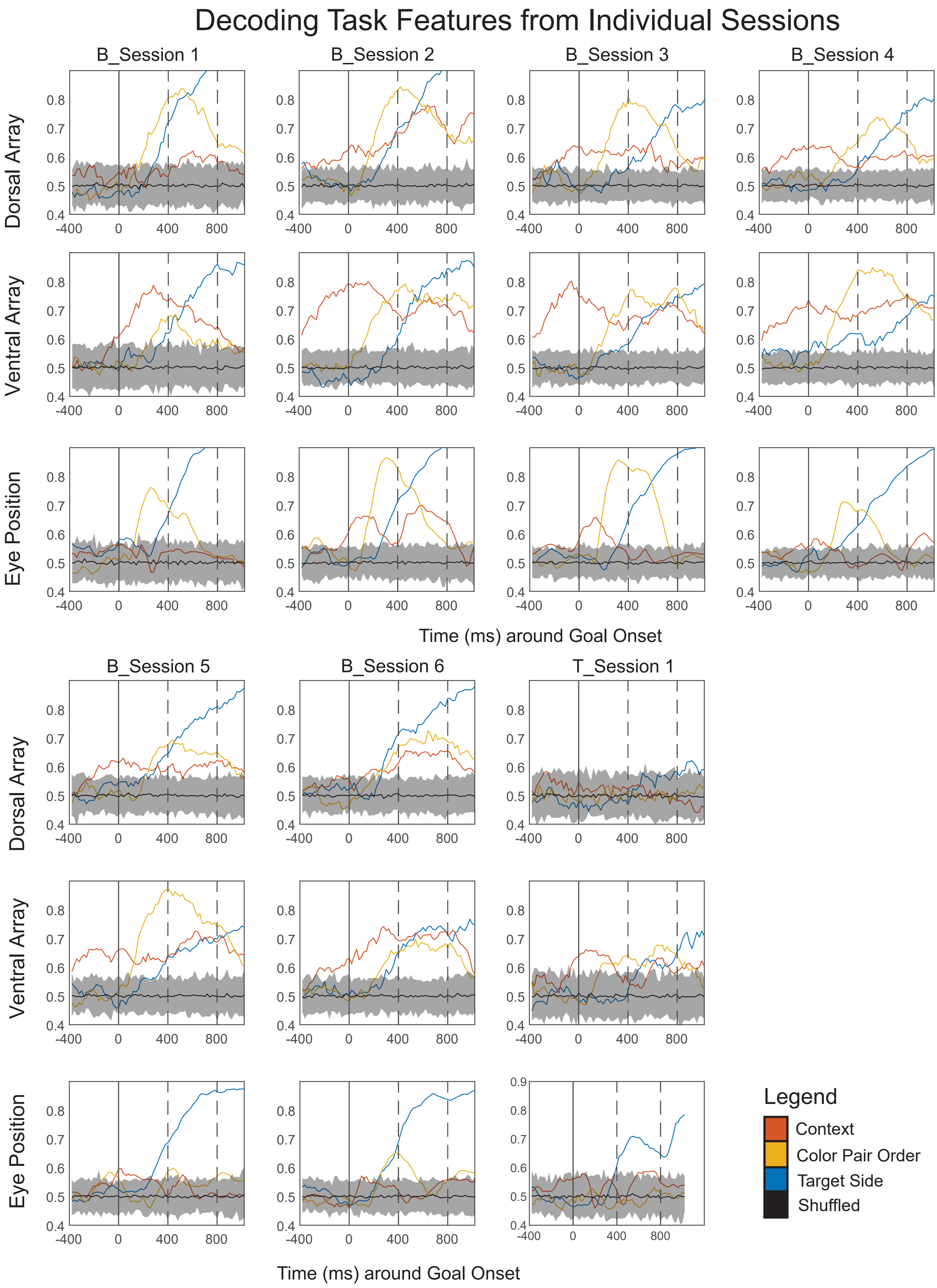

### Supplementary Figure 4

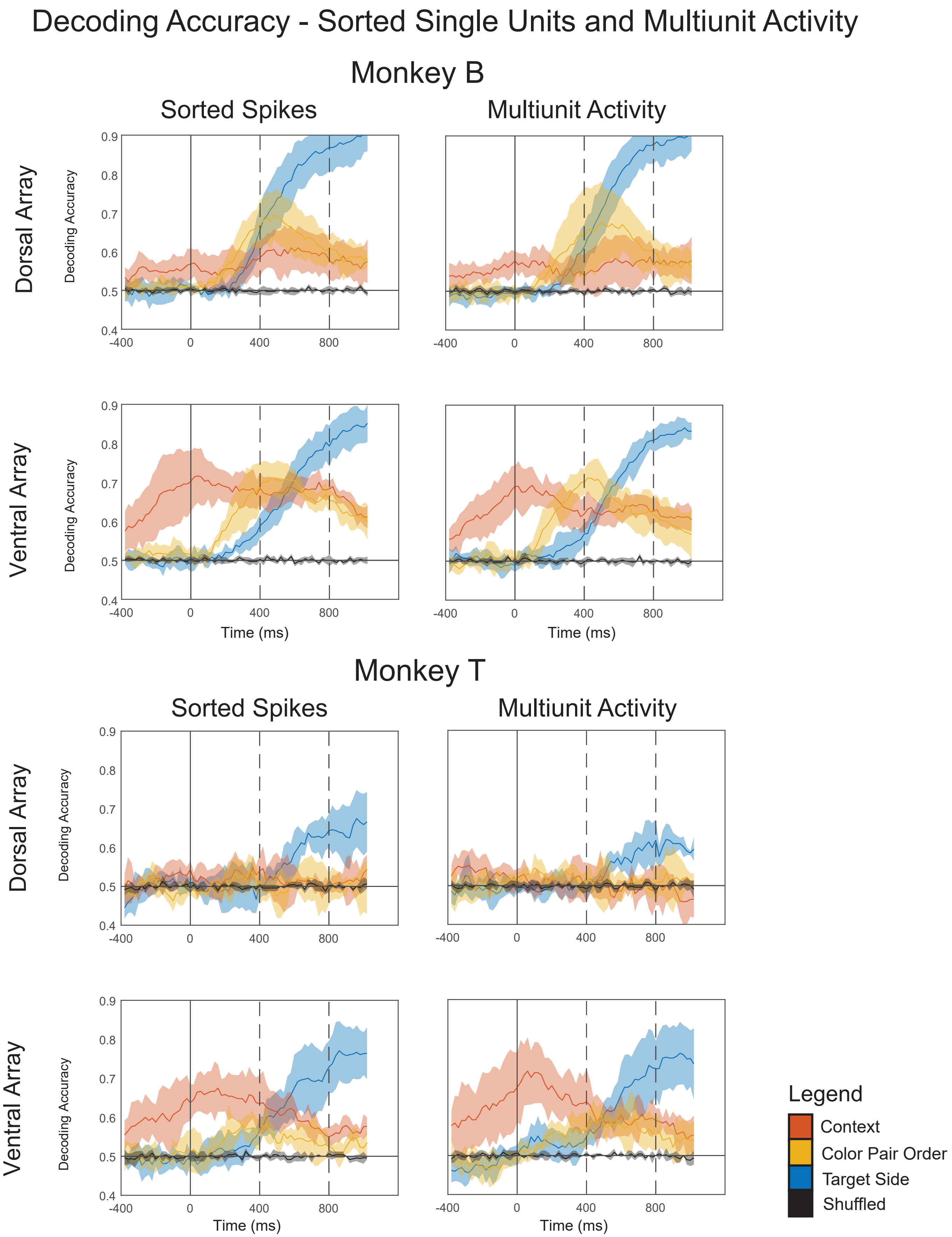

### Supplementary Figure 5

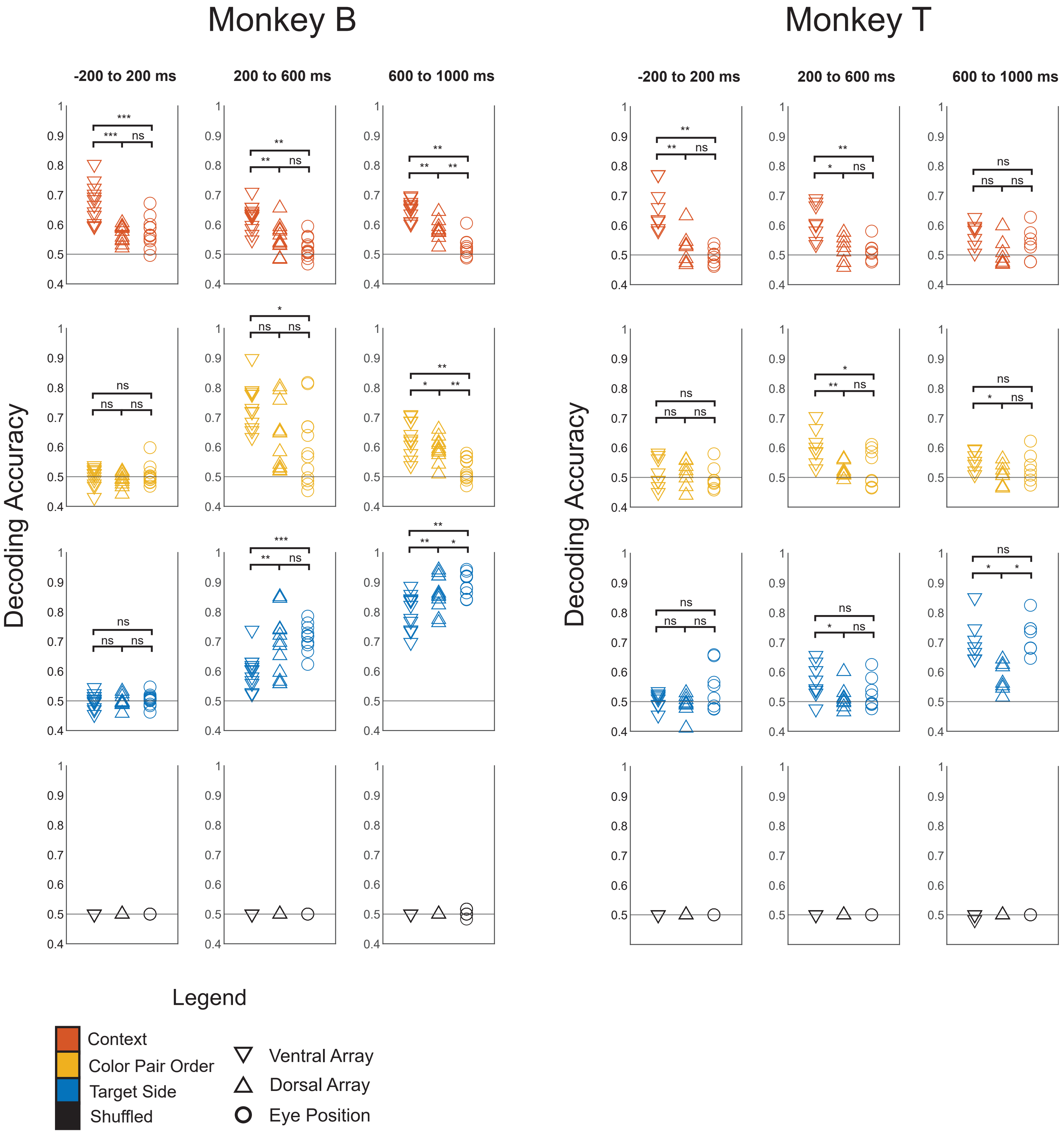

### Supplementary Figure 6

#### Dynamic Decoding - Monkey B

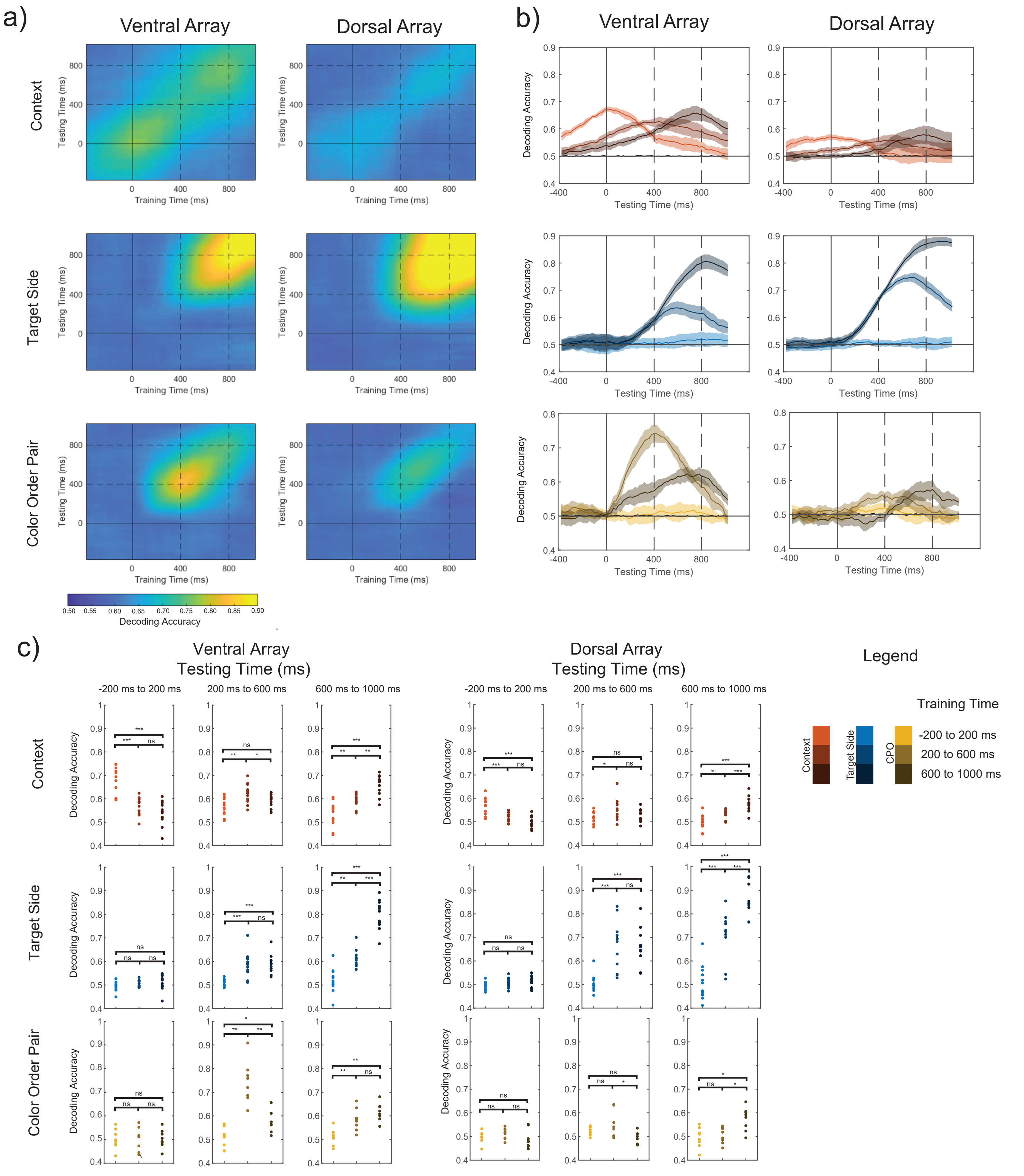

### Supplementary Figure 7

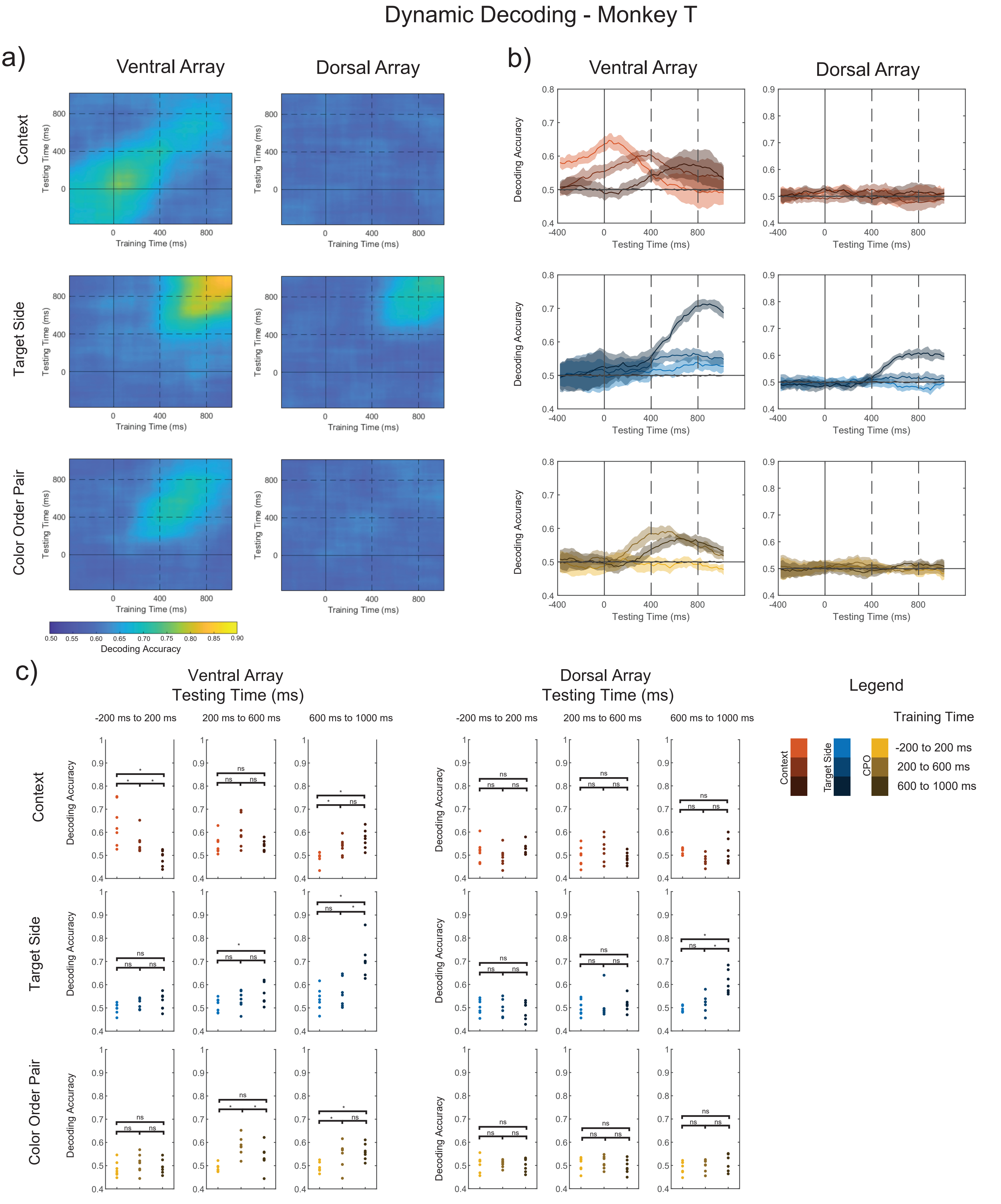
